# Interaural level difference sensitivity in neonatally deafened rats fitted with bilateral cochlear implants

**DOI:** 10.1101/2024.07.30.605756

**Authors:** Sarah Buchholz, Jan W. H. Schnupp, Susan Arndt, Nicole Roßkothen-Kuhl

## Abstract

Bilateral cochlear implant (CI) patients exhibit significant limitations in spatial hearing. Their ability to process interaural time differences (ITDs) is often impaired, while their ability to process interaural level differences (ILDs) remains comparatively good. Clinical studies aiming to identify the causes of these limitations are often plagued by confounds and ethical limitations. Recent behavioral work suggests that rats may be a good animal model for studying binaural hearing under neuroprosthetic stimulation, as rats develop excellent ITD sensitivity when provided with suitable CI stimulation. However, their ability to use ILDs has not yet been characterized. Objective of this study is to address this knowledge gap. Neontally deafened rats were bilaterally fitted with CIs, and trained to lateralize binaural stimuli according to ILD. Their behavioral ILD thresholds were measured at pulse rates from 50 to 2400 pps. CI rats exhibited high sensitivity to ILDs with thresholds of a few dB at all tested pulse rates. We conclude that early deafened rats develop good sensitivity, not only to ITDs but also to ILDs, if provided with appropriate CI stimulation. Their generally good performance, in line with expectations from other mammalian species, validates rats as an excellent model for research on binaural auditory prostheses.

## Introduction

Spatial hearing is a fundamental aspect of auditory perception. It enables us to precisely localize sound sources and makes it easier for us to parse acoustic environments with multiple sound sources. This remarkable ability depends to a large extent on our brain’s ability to process binaural directional cues, such as interaural time differences (ITDs) and interaural level differences (ILDs). In natural hearing, these occur when free-field sounds are presented from either side of the center line, resulting in differences in arrival time and in sound amplitude at each ear. Normally hearing human listeners are remarkably good at using those cues, and can detect ITDs as small as ∼10 µs [1] and ILDs as small as ∼0.5 dB [2-4]. Similarly small ITD detection thresholds can be found in a variety of other animals, although there are some species differences. While normally hearing ferrets and rats show ITD thresholds around 40-50 µs [5; 6], gerbils and guinea pigs can reach thresholds closer to humans with about ∼20 µs [7; 8]. ILD thresholds are similarly comparable across species, ranging from 0.5 dB in humans to 0.8 dB in gerbils, 1.3 dB in ferrets, 2.2 dB in rats, and 3 dB in guinea pigs [2; 3; 5; 7-9].

However, for severely-to-profoundly deaf patients who are dependent on bilateral cochlear implants (biCI) for their perception of the auditory world, binaural cue discrimination thresholds are often significantly impaired, especially for ITDs. This likely also impacts their ability to engage in other auditory scene analysis tasks. For example, biCI patients have difficulties understanding speech in noise, and they perform poorly in binaural tasks such as spatial release from masking or auditory scene analysis [10; 11]. These deficits in human cochlear implant (CI) patients are thought to be mostly due to impaired ITD sensitivity [12; 13], while ILD sensitivity often remains quite good [14-16], with some patients exhibiting ILD discrimination performance that is similar to that seen in normally hearing controls (in as far as ILDs of binaural acoustic and prosthetic stimuli can be directly compared).

What limits binaural cue sensitivity in biCI patients, and whether, or how, it might be improved with different treatment or sound processing strategies, remains an area of active investigation. The impaired ITD sensitivity often seen in biCI patients, which appears to be particularly pronounced in early deafened ones, may be due to a variety of factors that have the potential to interact. At present it remains extremely difficult to disambiguate the effects of suboptimal binaural coding strategies from possible critical period effects, or from the likely desensitizing effects that months of exposure to clinical processors may have which, as a general rule, do not deliver reliable ITD information in the timing of their electrical pulse trains. The clinical needs and the often quite varied clinical histories of patients make it impossible to recruit the large, homogenous patient cohorts and to subject them to the very specific treatment, training and testing regimes that would make it possible to isolate the many possible confounding factors in studies on human volunteers. These limitations associated with clinical research on human patients have motivated our group to turn to studies in rats, in which the potential confounds of an animals’ biological history, experience, and stimulation parameters can be precisely controlled. Rats already turned out to be a promising choice given that they are easy to train in behavioral binaural cue lateralization tasks [6] and highly suitable for electrophysiological investigations of binaural cue sensitivity [17-22]. In addition, the extremely encouraging results from Buck et al. [23] and Rosskohten-Kuhl et al. [24] showed, that neonatally deafened (ND), young adult cochlear implanted rats were able to learn with only minimal training to lateralize CI pulse timing ITDs as small as ∼50 μs for pulse rates up to 900 pps, exhibiting an ITD discrimination performance that is comparable to normally hearing peers and substantially better than that observed in comparable human patient cohorts [6]. These results are exciting because they open up avenues for further research into improved CI stimulation strategies or treatment regimes that should lead to substantially better outcomes for biCI patients. However, to build on those results, much work needs to be done, firstly to further validate the biCI rat as a good model for human prosthetic binaural processing by demonstrating that their ILD processing too is comparable to what is seen in humans.

In this study, we are taking a step towards this goal by addressing the question whether ND rats with biCI can develop good ILD sensitivity despite their deafness throughout their infancy, and, if so, whether this varies with pulse rate and is comparable to that of human biCI patients. We document, for the first time, the behavioral sensitivity to ILDs in a cohort of ND rats, which were bilaterally implanted with CIs in young adulthood and trained to lateralize binaural CI pulse trains in a 2-alternative forced choice (2AFC) procedure with positive reinforcement. The animals were then tested with synchronized binaural pulse trains of varying ILDs and pulse rates to generate psychometric functions to document their sensitivity to electric ILDs in considerable detail.

## Material and methods

All procedures involving experimental animals reported here were performed under license issued by the Regierungspräsidium Freiburg (#35-9185.81/G-17/124, #35-9185.81/G-22/067). We confirm that all of our methods were performed in accordance with the relevant guidelines and regulations and that our study is reported in accordance with the ARRIVE guidelines. A total of 24 female Wistar rats were used in this study. All rats underwent neonatal deafening, acoustic and electric auditory brainstem response (aABR/eABR) recordings, bilateral cochlear implantation and behavioral training as described in Rosskothen-Kuhl et al., Buck et al., Schnupp et al. [23-25] and briefly below.

### Neonatally deafening and cochlear implantation

All rats in this study were neonatally deafened by daily intraperitoneal injections of kanamycin (400 mg/kg) from postnatal day 8 to 20 inclusively as described in Rosskothen-Kuhl et al. [24]. Kanamycin is an ototoxic antibiotic known to cause widespread death of inner and outer hair cells [26; 27] Severe to profound hearing loss (>80 dB) was confrimed by the loss of Preyer’s reflex [28] and the absence of ABRs to broadband click stimuli.

The animals were raised to young adulthood (postnatal week 10-14) and then implanted simultaneously with biCIs under ketamine (80 mg/kg) and xylazine (12 mg/kg) anesthesia. We implanted two to three electrodes of an electrode array either from PEIRA (animal array ST08.45, Cochlear Ltd, Peira, Beerse, Belgium) or from MED-EL (3-patch animal arrays, Medical Electronics, Innsbruck, Austria). The electrodes were inserted via a cochleostomy over the middle turn, and their lead wires were connected to a Hirose connector which was fixed with dental acrylic on the vertex of the animal’s cranium to allow direct electrical stimulation of the cochlea. The correct function of the CIs was confirmed using eABRs.

### Electric stimulation

The electric stimuli used to examine the animals’ eABR and behavioral ILD sensitivity were generated using a Tucker-Davis Technology (TDT, Alachua, FL) IZ2H programmable constant current stimulator at a sample rate of 48,828.125 Hz, and delivered directly to the intracochlear electrodes via a custom made cable connecting the IZ2H to the head-mounted percutaneous connector just described. The cable included a slip-ring connector and was suspended above the animal’s head from a counterweighted balance arm that ensured that the animal could move freely in the behavior box without the cable getting tangled or impeding the animal’s movement.

One of the tip CI electrodes served as the stimulating electrode, the adjacent electrode as ground electrode. All electrical intracochlear stimulation used biphasic current pulses similar to those used in clinical devices (duty cycle: 40.96 µs positive, 40.96 µs at zero, 40.96 µs negative), with peak amplitudes of up to 300 μA, depending on eABR thresholds and informally assessed behavioral comfort levels (rats will scratch their ears frequently, startle or show other signs of discomfort if stimuli are too intense). For behavioral training, all neonatally deafened, cochlear implanted (NDCI) rats were stimulated with an average binaural level (ABL) of ∼2-10 dB above these thresholds depending on the pulse rate. Careful observation of the animals’ behavior during spontaneous presentations of test stimuli was used to confirm that stimulus levels were high enough to be reliably detected, but not so high as to cause discomfort. Behavioral stimuli from the TDT IZ2H were delivered directly to the animal through a custom-built head connector that was connected and disconnected before and after each training session. For full details on the electric stimuli and stimulation setup see Rosskothen-Kuhl et al., Buck et al., Schnupp et al. [23-25].

### Psychoacoustic training and testing

After implantation, the rats were trained in a 2AFC stimulus lateralization task in a custom-made behavioral setup [23-25]. The setup is illustrated schematically in Figure 1A. It consists of a cage with three water spouts on one side. The rats were trained to initiate trails by licking the center spout first, which triggered bilateral CI pulse trains. The animals then had to choose between the left and right response spout according to the stimulation. Triggering the correct spout was rewarded with water, while triggering the wrong spout resulted in a short (∼10-15 s) time-out.

**Figure 1.**
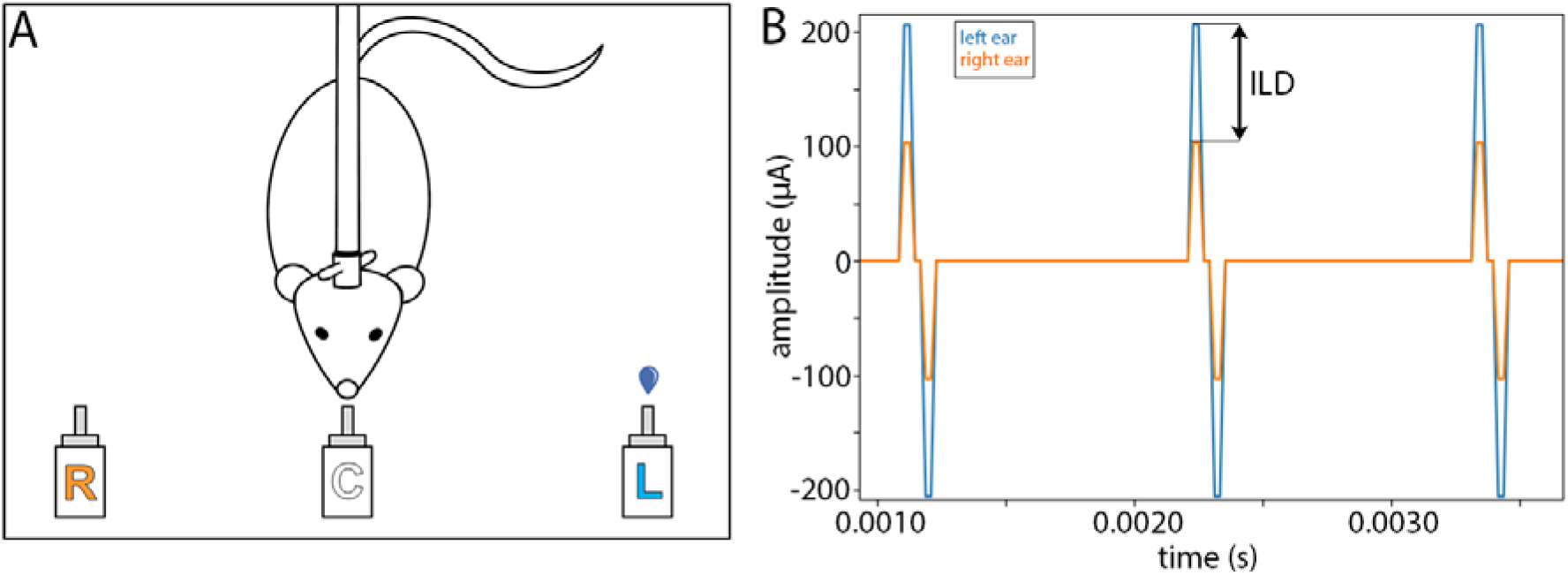
(A) Neonatally deafened, cochlear implant (CI) supplied rat performs a two-alternative forced choice sound lateralization task in our custom-build behavioral setup. The rat licks the center spout (C) and thus triggers the presentation of a bilateral pulse train via the CIs. According to the interaural level difference (ILD) on the pulse train, the rat must give a response at the left (L) or right (R) water spout to be rewarded. In the example shown in (B) the rat would need to respond on the left spout to receive drinking water as positive reinforcement. An incorrect response would lead to a timeout. (B) Example of a binaural electric stimulus pulse train with an ILD of 6 dB favoring the left ear.

Prior to testing for ILD sensitivity, the rats took part in another stimulus lateralization study performed in the same setup which will be reported elsewhere. Consequently they were already familiar with the idea of initiating trials by licking the central spout and using cues from bilateral CI stimulation to guide their behavioral choices when we began training them on lateralizing stimuli on the basis of ILD cues alone. The pulse trains were up to 5 s long, but animals were free to make a choice as soon as they wished after stimulus onset, and the stimulus was terminated as soon as the animal touched one of the response spouts. On average, animals responded within approximately 2 s after stimulus onset. After about five weeks of training and testing, the rats were tested for their ILD sensitivity at 900 pps by presenting unmodulated, rectangular pulse trains with ITD = 0 µs and different ILD values ±{0.5, 1, 2, 3, 4, 5, 6} dB. Figure 1B shows the current waveforms sent to each ear for a 6 dB ILD, 900 pps pulse train stimulus. Throughout the ILD training and testing, the animals only received precisely synchronized (zero ITD) input on both ears, which differed only in the amplitude of the pulse train, i.e. the ILD. Prior to testing, the ABL at which they were tested was confirmed by carefully observing the animal’s reaction to short, unexpected stimuli.

All animals had to lateralize stimuli for which stimulus amplitudes in both ears were always 2-10 dB above the eABR and behavioral threshold. To exclude the possibility that the animals used monaural level cues instead of binaural level comparisons to solve the task, some of the rats were tested with the same ILDs at different ABLs, which confirmed that the animals’ performance collapsed with near-threshold or unilaterally suprathreshold stimuli. Five of the NDCI rats received additional testing for their ILD sensitivity at 50, 300, 1800, and 2400 pps under the same conditions. Since higher pulse rates at equal amplitudes are commonly perceived as louder, the appropriate ABLs were determined separately for each pulse rate tested. ILD sensitivity testing for a given pps extended over 4-8 sessions. The animals typically performed about 200 trials in each session, yielding a total of approximately 800 trials for each of the ILD psychometric curves reported below.

## Data analysis

To determine behavioral ILD sensitivity, an animal’s proportion of “right” responses (pR) was fit as a function of stimulus ILD using either linear or cumulative Gaussian sigmoid functions (Figs. 2, 4), for comparison see Rosskothen-Kuhl et al. [24]. Briefly, the psychometric data were fit with three different models:

**Figure 2.**
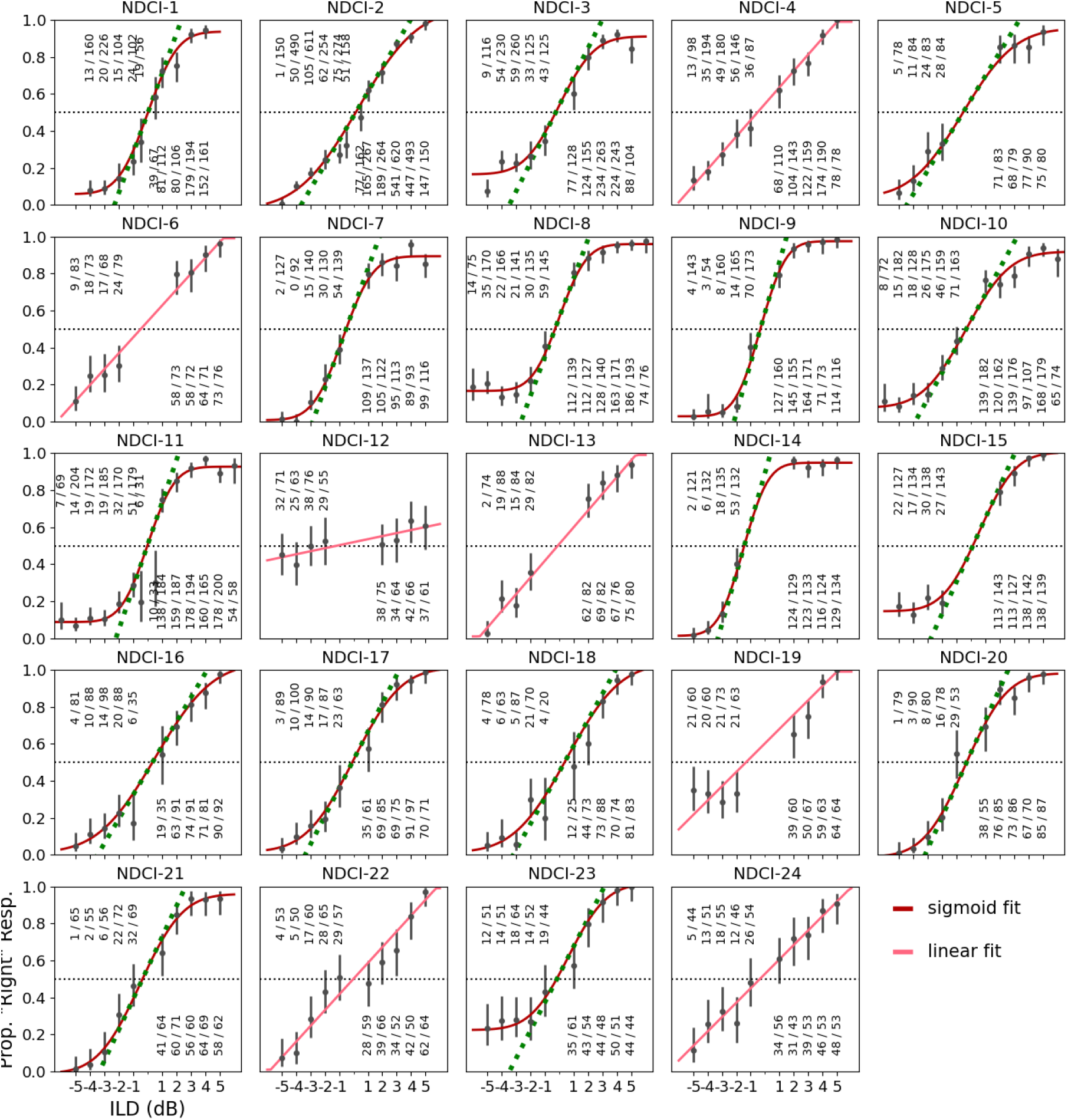
Psychometric functions for interaural level difference (ILD) lateralization for 24 neonatally deafened (ND) rats with bilateral cochlear implants (CIs). X-axis: ILD in dB, with negative values representing stimuli louder on the left. Y-axis: proportion of trials in which the animal responded on the right side. The fractions above each tested ILD value show the observed number of ‘right’ responses over the total number of trials. Error bars: 95% Wilson confidence intervals for the underlying probability of a ‘right’ response. Continuous red lines show the fitted psychometric models. If the best fit was a sigmoid model it is shown in dark red, and if it was a linear model it is pictured in light red as shown in the figure legend. The green dotted line superimposed on the sigmoid fits show the slope of the fitted psychometric function at ILD=0 dB. The horizontal black dotted lines demarcate 50% ‘right’ responses, the performance one would expect if the animals were insensitive to ILDs and responded simply with unbiased guessing.

a modified Probit regression (sigmoid) model

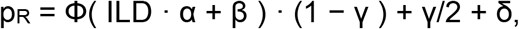

a bound linear model

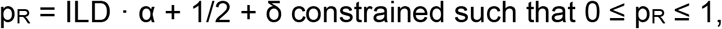

or a null model

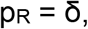

where ILD is the difference (right - left) in stimulus pulse amplitude in dB, α captures the animal’s sensitivity to ILD, β a possible “ear bias” (i.e., 0 ILD may be heard off-center), δ captures a possible “spout bias” (i.e., when an animal guesses, it may have an idiosyncratic preference for one side), and γ is a lapse rate that captures the proportion of times the animal makes errors due to inattention or exploratory behavior, even though the stimulus should be easy to discriminate. A deviance test was used to determine which of the three possible models best fit the behavioral data. Once the best fitting model was obtained, the animals’ “just noticeable difference” (JND) threshold was calculated by using the model equation to compute the change in ILD in dB required to raise the animal’s pR from 50 % to 75 % (equivalent to a 75 % correct threshold after correcting for possible spout biases). In addition, the slope of the fitted psychometric curve at ILD = 0 dB was calculated in units of % change in the animal’s preference for choosing the correct water spout per dB increase in the value of the stimulus ILD. The slope together with the JND served as measures of the animal’s behavioral sensitivity to ILDs.

## Results

Twenty-four ND rats received chronic bilateral CIs in early adulthood (postnatal weeks 10–14) and were trained on a 2AFC sound lateralization task and subsequently tested for their ILD sensitivity at a typical clinical pulse rate of 900 pps as just described.

Figure 2 shows the behavioral ILD performance of each rat in the form of psychometric curves. As might be expected, the behavioral performance varied slightly from animal to animal, but overall a very good ILD sensitivity was observed for 23 out of the 24 rats tested at 900 pps, as evidenced by the good fit of steep sigmoidal or linear psychometric function fits to the data, and low error rates for absolute ILDs >2dB. The only exception is rat NDCI-12, which performed near-chance even at large ILDs. However, not in a single case did the null model produce the best fit to the data, which indicates that every rat in the sample of 24 animals exhibited sensitivity to ILD.

To quantify the behavioral ILD sensitivity of each rat, we calculated the slopes of the psychometric curves at ILD = 0 dB, and to facilitate comparisons with other studies we also computed a 75 % correct threshold (sometimes described as a “just noticeable difference” (JND) value) from the fitted psychometric curves. The distribution of the observed behavioral ILD sensitivities, psychometric slopes and JNDs for all 24 NDCI rats, are summarized in Figure 3. The median slope across all animals was 14.5 %/dB, and the median JND for ILDs was 1.715 dB.

**Figure 3.**
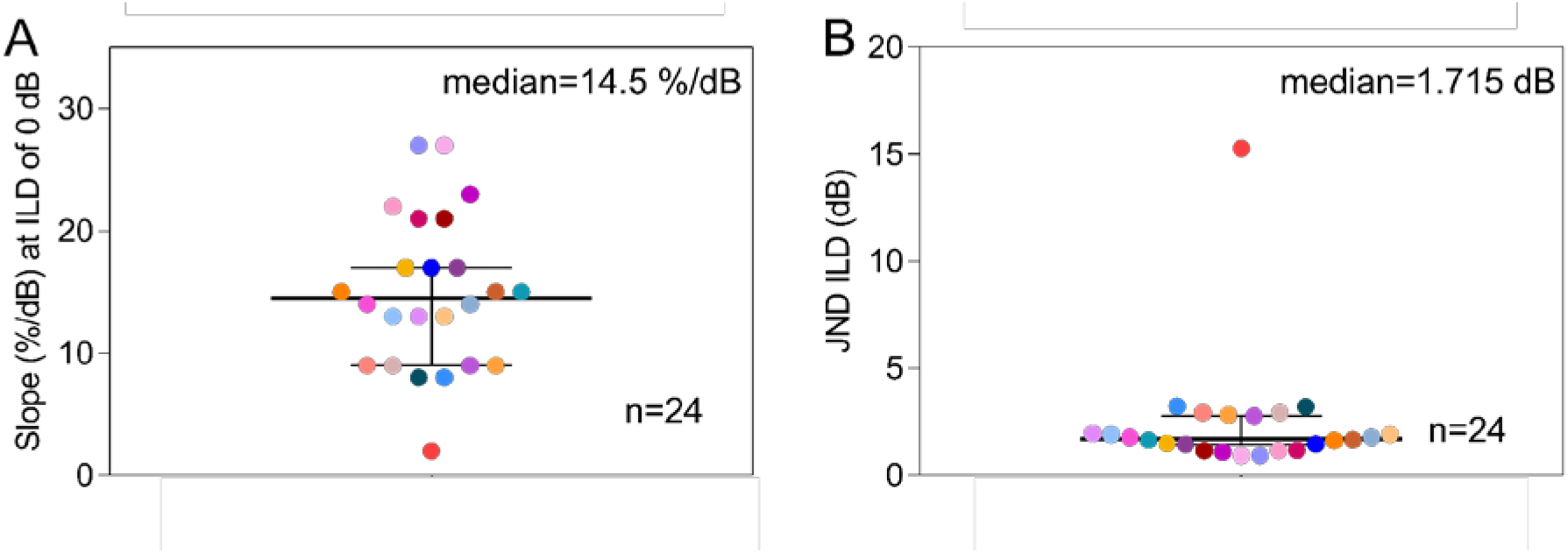
Swarm plots showing the distributions of the interaural level difference (ILD) sensitivity values observed across a cohort of 24 neonatally deafened cochlear implanted (NDCI) rats. (A) Distribution of ILD sensitivity quantified by the slope of the psychometric curve around zero ILD (cp. to Fig. 2, green dashed line or red line). (B) Distribution of ILD sensitivity quantified as just noticeable difference (JND). Each rat is represented by a different color.

**Figure 4.**
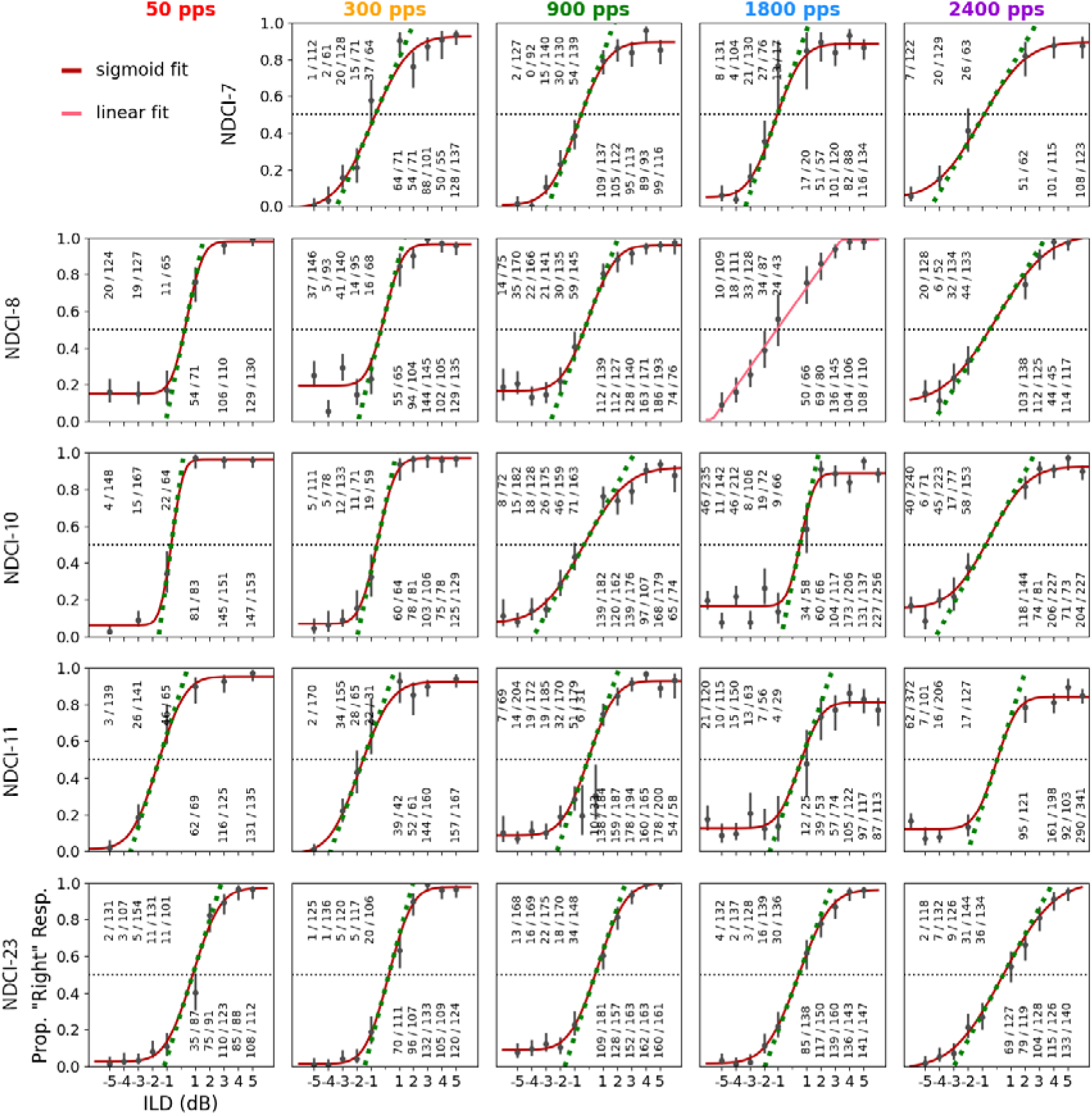
Psychometric functions of the interaural level difference (ILD) lateralization performance for five neonatally deafened (ND) rats with bilateral cochlear implants (CIs) for five different pulse rates. The rows show the five different rats (NDCI-7, NDCI-8, NDCI-10, NDCI-11, NDCI-23) and the columns represent the five different pulse rates (50, 300, 900, 1800 and 2400 pps) used for testing. Details are as for Fig. 2.

Additionally, five out of 24 NDCI rats were tested on their ILD sensitivity at other pulse rates. All five NDCI-rats were tested at 300, 1800, and 2400 pps, and four of the five animals were also tested at 50 pps. The psychometric data from these five rats are shown in Figure 4. All rats showed very good ILD sensitivity, with steep slopes and low error rates at large absolute ILDs, at all pulse rates tested. The slopes and JND values computed from these psychometric functions are summarized in Figure 5. The different pulse rates are shown in different colors and the five rats can be identified by different symbols as described in the figure legend.

**Figure 5.**
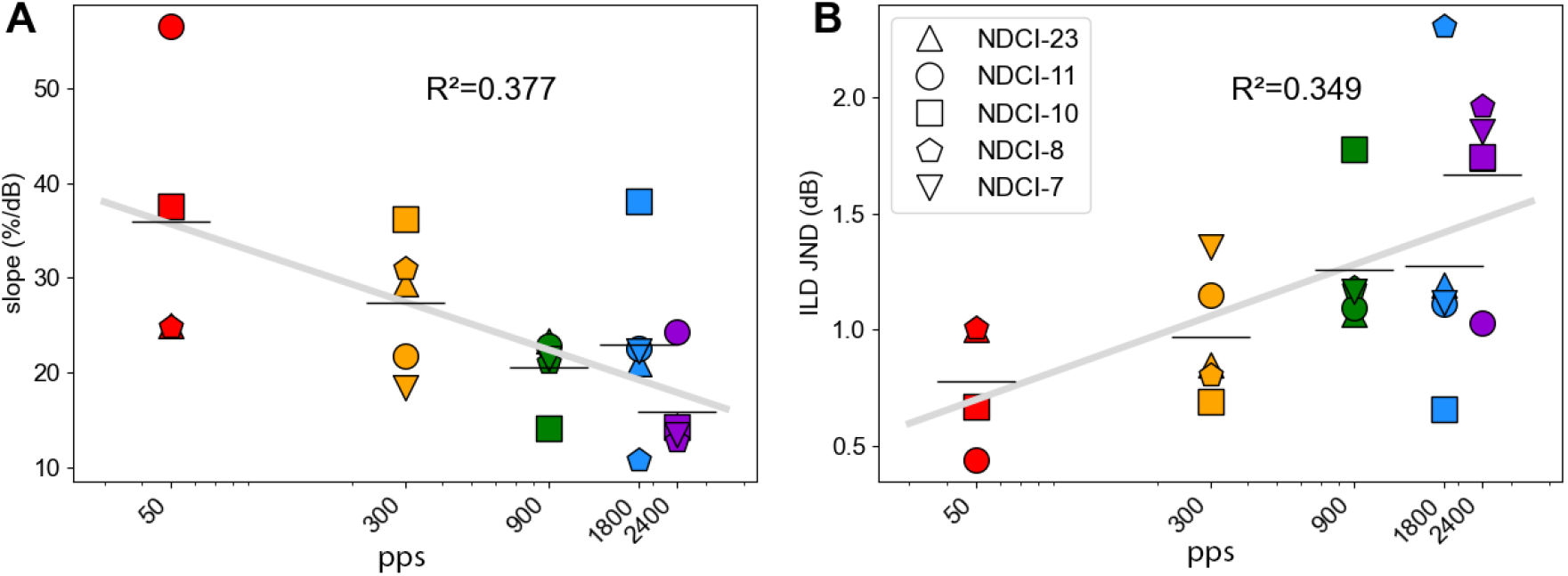
Relation between pulse rates (pps) and interaural level difference (ILD) sensitivity in electric hearing. The colored symbols show the ILD sensitivity for each of the five neonatally deafened, cochlear implanted (NDCI) rats tested for five different pulse rates (50, 300, 900, 1800, and 2400 pps). The five rats are shown as different symbols as indicated in the legend. The horizontal black lines show the mean ILD sensitivity value across animals for each pulse rate. Thick gray line: linear regression line with the coefficient of determination (R^2^) shown. (A) ILD sensitivity quantified by the slope of the psychometric functions (Fig. 2, green dashed line or light red line). (B) Estimated ILD just noticeable difference (JND) values as a function of pps.

Across the five NDCI rats tested, ILD sensitivity tended to be dependent on the pulse rate. The average slope decreased with increasing pulse rate, and conversely the JND ILD increased with increasing pulse rate (Fig. 5). This becomes particularly clear when comparing mean JND ILDs. For example, the mean JND at 300 pps (1 dB) was only about half as large as that at 2400 pps (1.7 dB). To determine possible dependencies between the variables pulse rate and slope or pulse rate and JND, we carried out a linear regression of log (pps) against ILD sensitivity (gray lines in Fig. 5). The R-squared value for the slopes was 0.377 and 0.349 for the JNDs, and the regression slopes were significantly different from zero (p=0.001 for slopes, *p*=0.002 for JND). Our data thus indicate a statistically significant trend for ILD sensitivity to decline with increasing pulse rates.

## Discussion

In this study, we set out to measure and describe the behavioral sensitivity of ND rats to ILDs in pulsatile electric stimuli delivered binaurally via neuroprosthetic intracochlear electrodes. As explained in the introduction, rats have turned out to be a very useful model species for investigating ITD sensitivity under biCI stimulation, as they exhibit remarkably low behavioral [23-25] and physiological [20; 24] ITD thresholds even after neonatal deafening and over a wide range of pulse rates. The use of this animal model allows us to circumvent the many confounds that patient needs and clinical practice issues introduce into studies on human volunteers, making it easier to investigate factors that determine binaural hearing outcomes following bilateral cochlear implantation. To complete the characterization of rats as a model species of prosthetic binaural hearing, it is pertinent to document their sensitivity to ILDs.

### Acoustic ILD sensitivity across mammalian species

Acoustic ILD sensitivity is fairly comparable across mammalian species. We note that acoustic ILD JNDs reported for mammals tend to lie in the range from just below 1 to about 3 dB; 3 dB for normally hearing rats presented with pure tones over loudspeakers [29], 1.3 dB for ferrets [5], 0.8 dB for gerbils [7], 3 dB for guinea pigs [8] and values in a similar range of 0.5 to 2.5 dB for humans [2; 3; 14; 30]. None of this is terribly surprising given that much homology is thought to exist between mammalian species in the auditory brainstem pathways responsible for computing binaural cues [30; 31] even if there may be differences in the extent to which individual species rely on ITD, ILD or both. Two important auditory brainstem regions for ILD processing are the lateral superior olive (LSO) and the inferior colliculus (IC). The LSO is well-developed in all terrestrial mammals studied to date [32-34] and similar anatomy across mammalian species is also found in the IC, a central hub for processing binaural input from both ears [35-37]. Therefore, from an anatomical point of view, there is no reason to assume ILD insensitivity in mammals although it remains to be seen how early deafness affects the ILD perception with auditory prostheses in mammals such as the rat.

### ILD sensitivity in the deaf auditory system

How disturbances of auditory input affect the development of the brain’s circuitry for ILD sensitivity has, to the best of our knowledge, been little studied to date. A recent study by Hubka et al. [38] demonstrates that neurons of the auditory cortex of bilaterally congenital deafened cats, supplied with CIs, exhibit unaltered tuning in comparison to normal hearing cats, although the modulation amplitude of ILD functions and the firing rate were reduced. However, in unilaterally deafened cats with bilateral CIs, ILD representation becomes inconsistent and is dominated by the hearing ear, which seems to be due to subcortical reorganization caused by unilateral auditory deprivation. Polley et al. [39] also looked at the effect of unilateral deafness, but in the context of reversible unilateral conductive hearing loss at different time points in the developing primary auditory cortex of mice. They show that the binaural response properties of the auditory cortex are rather mature at the onset of hearing with exception of ipsilateral ILD sensitivity. Furthermore, reversible unilateral conductive hearing loss at P16 disrupts sensitivity to ipsilateral ILDs, implying significant changes during the early auditory period. Nevertheless, regarding early bilateral hearing loss one would not necessarily expect a severe disruption of ILD sensitivity if hearing is later restored, given that as for example Buck et al. [20] demonstrated that even the hearing inexperienced brain is exquisitely sensitive to ITDs, suggesting that high quality auditory experience early in life may not be necessary for functional binaural circuitry to emerge in. This hypothesis is supported by the study of Couchman et al. [40], who performed whole cell patch electrode recordings from LSO principal neurons in a mouse model of congenital deafness and found that inhibitory input to LSO develops normally. Furthermore, using a circuit model, they were able to show that a functional ILD circuit appears to be maintained during congenital deafness.

### ILD sensitivity in cochlear implant users

In agreement with these observations, an unaffected ILD sensitivity despite a lack of early hearing experience can also be observed in prelingually deafened CI patients [15; 41; 42]. Litovsky et al. [43] even studied patients with different onset of deafness, and found that patients who were early deafened did not show less robust ILD sensitivity than late deafened patients. The generally good ILD sensitivity in human CI patients for binaural stimuli presented via the audio inputs of the clinical devices ranges from 1 to 5 dB and thereby is largely comparable to normal hearing subjects with thresholds ranging from 0.5 to 2.5 dB, in as far as sensitivity to interaural electric versus acoustic stimulus intensity differences can be directly compared [2; 3; 14; 16; 44]. Meanwhile, the ILD sensitivity in human CI patients tested via loudspeaker seems to be slightly lower with ILD JND of in mean 3.8 dB as observed by Grantham et al. [45]. Overall, human CI patients are fairly ILD sensitive not only with research processors tested at low stimulation rates [41; 43; 46] but also for higher stimulation rates using their own, clinical CI processors [42; 45; 47]. Within the framework of this study we could now confirm this very good ILD sensitivity of biCI users by employing the animal model of NDCI rats. Our NDCI rats learned to lateralize ILDs of only a few dB in neuroprosthetic electrical stimuli despite the absence of early hearing experience (Fig. 2, and 3), with best ILD JNDs at about 1 dB (Fig. 3 B), a value fairly comparable to the thresholds observed in human CI patients [14; 16; 44].

Furthermore, the thresholds of our early deafened CI rats are fairly comparable to those of normal hearing rats (∼2.2 dB), tested behaviorally with acoustic broadband noise stimuli via headphones fixed on their head [9]. This is an interesting result considering the fact that electric and acoustic hearing are not directly comparable. Indeed, comparisons between electrical and acoustic ILD values are not at all straightforward, not only because the units of amplitude differ, but also because electrical stimulation of the inner ear bypasses the non-linear outer hair cell amplifier and the numerous adaptive mechanisms at the inner hair cell-auditory nerve fiber synapses. These allow acoustic hearing to operate over an astonishing 120 dB dynamic range, while that of electric hearing is confined to a much less impressive electric dynamic range of 8-20 dB [48; 49]. ILD sensitivity measured when stimuli are presented to the audio inputs of a CI device will therefore depend on how the device processor maps the acoustic inputs onto electric stimulus outputs just as much as on how sensitive the CI patients are to electric stimulus amplitude ILDs. Thus, there are a multitude of both technical and biological factors that complicate comparisons between the sensitivity to ILDs in stimuli presented acoustically to normally hearing individuals, to the audio inputs of clinical devices, or in the amplitude of electrical pulse trains delivered via intracochlear electrodes.

### ILD sensitivity across pulse rates

In addition to a very good ILD sensitivity at a clinically stimulation rate of 900 pps, we identified good ILD sensitivities at different pulse rates for five tested NDCI rats. Interestingly, the ILD sensitivity tended to decrease with higher pulse rates. As shown in Figure 5B, the best ILD JND was found at 50 pps, while the ILD JND for 2400 pps was more than twice as high. A similar negative dependence on increasing pulse rates has previously been described for ITD sensitivity in acoustic [6] and electrical binaural hearing [23]. Normally hearing rats showed a strong decrease in ITD sensitivity from 50 to 4800 pps regardless of the shape of the envelope. Comparable observations were made in NDCI rats, although the best ITD JND was observed at 300 pps for stimuli with a rectangular envelope [23]. The exact mechanisms underlying the dependence of binaural sensitivity on pulse rate remain to be determined. However, it is possible that, at low pulse rates, the auditory pathway may still process cues conveyed in each pulse separately and to an extent independently, providing the brain with multiple opportunities in each pulse train to estimate binaural cue values and to combine these estimates to yield higher sensitivity. But the possibility to do this may decline as later pulses in a fast train are increasingly poorly resolved and partly suppressed by adaptation.

### Rats as a model species of prosthetic binaural hearing

The ILD sensitivity documented here for our NDCI rats appears to be in line with what might be expected in the light of previous human CI studies. These findings additionally, with rats’ high sensitivity to ITDs as previously [23-25] reported makes the NDCI rat an excellent model for human binaural hearing under electric stimulation. Thereby, the unbeatable advantage of our animal model compared to human patients is complete control of all test parameters, like etiology and technical factors. Since most standard clinical sound processors produce pulsatile electrical stimulation with fixed pulse rates and independent pulse timing, the ITDs delivered to the patient’s ears are random and unrelated to the binaural cues of different sound source directions. Such devices may still encode ITDs in the envelopes of pulse trains delivered to the left and right ears, respectively [50], but the electrically stimulated auditory pathway may find it hard to ignore random pulse timing ITDs while paying attention of potentially informative envelope ITDs. However, in our NDCI rats we can present microsecond precise temporal information via the CIs from the outset. Under this stimulation, NDCI rats are not only able to discriminate ITDs of only 50 µs, comparable to normal hearing littermates [6; 24], but also, as shown for the first time in this study, ILDs of a few dB with thresholds comparable to those of normal hearing rats and human CI patients [9; 14; 44].

## Conclusion

Our data have shown that NDCI rats readily learn to lateralize ILDs of binaural pulse trains at different pulse rates, with ILD discrimination thresholds that are broadly in line with what might be expected based on previous behavioral studies on humans and a variety of other mammals. Taken together with the recent observation that NDCI rats also easily learn to lateralize even very small ITDs, these results evidence that rats are a highly suitable animal model for the study of binaural processing under neuroprosthetic stimulation with the advantage to overcome limitations of human volunteers such as varied clinical histories including different hearing experience.

## Acknowledgements

The work leading to this publication was supported by the German Academic Exchange Service (DAAD) with funds from the German Federal Ministry of Education and Research (BMBF) and the People Programme (Marie Curie Actions) of the European Union’s Seventh Framework Programme (FP7/2007-2013) under REA grant agreement n° 605728 (P.R.I.M.E. – Postdoctoral Researchers International Mobility Experience), the Research Commission of the Medical Faculty of the Medical Center at the University of Freiburg, and the charity “Taube Kinder lernen hören e. V.“, MED-EL Medical Electronics, Innsbruck, Austria (Research Agreement PVFR2019/2), as well as grants from the Hong Kong General Research Fund (11100219 & 11103823), the Shenzhen Science Technology and Innovation Committee (JCYJ20180307124024360) and the Martin Lee Centre for Innovations in Hearing Health at Macquarie University. Thirty-two cochlear implant animal arrays were kindly provided by MED-EL Medical Electronics, Innsbruck, Austria (Research Agreement PVFR2019/2). The authors would also like to thank Theresa Preyer and Henrike Budig for their assistance in collecting data.

## Author contributions

JS, NRK, and SB designed research; NRK and SB performed the surgeries; SB collected the data; JS and SB developed the analysis pipeline; SB, JS, and NRK analyzed the data; SB, NRK, JS and SA discussed the data; SB, NRK and JS wrote the paper.

## Data availability

All data as well as the analysis code used to generate all the figures and statistical results included in this manuscript are available from the corresponding author on reasonable request. All data generated or analyzed during this study are included in this published article.

## Additional information

The authors declare they have no competing interests.

## Notes

### Competing Interest Statement

The authors have declared no competing interest.

